# Molecular epidemiology of *Ascaris lumbricoides* following multiple rounds of community-wide treatment: who infects whom?

**DOI:** 10.1101/2024.12.20.629811

**Authors:** Toby Landeryou, Rosie Maddren, Jack Hearn, Mahlet Belachew, Santiago Rayment Gomez, Ewnetu Firdawek Liyew, Kathryn Forbes, Birhan Mengistu, Scott Lawton, Jude Eze, Geremew Tasew, Ufaysa Angulo, Roy Anderson

## Abstract

Control and elimination of the parasite *Ascaris lumbricoides* relies on mass administration using a limited number of anti-helminthics. Whilst these programs have reduced the infection intensity and prevalence within many endemic regions transmission is poorly understood, with reinfection commonly occurring following cessation of treatment. Here, we utilise genomic data to understand parasite transmission within and between households in a community and the genomic impact of repeated MDA. We sequenced 54 whole-genomes from individuals in a longitudinal cohort epidemiological study of transmission and drug treatment extending over 6 years. We found that fine-scale population structure exists in spatially distinct clusters of infected individuals with reinfection occurring within or between geographically close households. This observation helps inform the policy of future control in low prevalence settings suggesting more targeted treatment of infection hotspots We found evidence of positive selection acting on members of gene families previously implicated in reduced drug efficacy but detect no high frequency of impactful variants. As efforts to eliminate *A. lumbricoides* intensify, our study provides a foundation for genomic surveillance to help identify both who infects whom and the impact of repeated drug treatment.

## Introduction

The soil-transmitted helminths (STH) are a group of intestinal parasites designated by the World Health Organisation (WHO) as the aetiological agents of a neglected tropical disease. There are an estimated 1.5 billion individuals infected with at least one intestinal nematode infection globally, cumulatively resulting in over five million disability-adjusted life years (DALY’s) ^1^. *Ascaris lumbricoides* is one of the most prevalent species of STH, with infection occurring through the ingestion of embryonated eggs from contaminated soil, food and/or water. Ascariasis is estimated to affect ∼819 million people globally ^2^, predominantly school age children (SAC) who on average harbour a greater number of parasites than adults. Treatment is delivered via mass drug administration (MDA) campaigns of the anthelminthic drugs albendazole and mebendazole within communities on an annual or bi-annual basis, dependent on the intensity of transmission ^3^.

Over the past three decades, these chemotherapeutic control programmes have been effective at reducing the prevalence and intensities of infection to low levels in many endemic regions (<u>Soil-transmitted helminthiasis | ESPEN</u>). Following these successes, the WHO set a goal to achieve elimination as a public health problem of *A. lumbricoides* (defined as <2% heavy or moderate intensity infections) in pre- and SAC in all 78 countries with endemic infection, and eliminating transmission in selected regions by 2030 ^4^. The currently recommended treatment strategy only targets SAC and pregnant women, however to reach these goals, some recently implemented deworming projects have expanded treatment strategies from targeting only school-based MDA (sMDA) to community-wide MDA (cMDA) ^5–7^. To achieve transmission, break a 2% prevalence must be achieved at a community and region-wide level, supported through mathematical models of STH transmission and control, and several large-scale epidemiological studies ^8–11^. Untreated infected individuals (untargeted adult population and untreated SAC), with high worm burdens act as reservoirs of infection for recently treated and STH-cleared individuals and hence community wide treatment is required to eliminate all transmission within defined populations ^12^. Furthermore, prior infection does not confer strong acquired immunity. Therefore, especially in areas with poor access to adequate water, sanitation and hygiene (WaSH) infrastructure, repeated re-infection is common which frequently results in rapid bounce-back of local infection after cessation of MDA ^10,13–15^. Where treatment coverage is not delivered at a consistently effective level to the target populations, a constant cycle of treatment and reinfection pertains.

The distribution of *A. lumbricoides* infection in endemic populations is over dispersed or aggregated which is well described by the negative binomial probability distribution, where most harbour none or a few worms and a few are infected by many ^16^. This aggregation is measured by k, the negative binomial parameter. As chemotherapeutic pressure increases over repeated rounds of MDA, the degree of worm aggregation also tends to increase ( very low k values ^17,18^) creating hotspots of infection in the community which are challenging to identify through routine surveys and therefore treat. Those heavily infected tend to be predisposed to infection

Accurately measuring changes in parasite population size and structure within these low prevalence communities is crucial to an understanding of the current and future impact of expanded MDA. Tracking how these few individuals with the remaining worms acquire infection and sustain infection in the wider community is key to the successful ‘end game’ STH control in regions that have had decades of MDA. The renainging pockets of infection may result due to many by many factors including systemic non-compliance of households to treatment, poor WaSH status, or social and cultural factors influencing exposure to infection^19^.

Establishing ‘who infects whom’ via molecular epidemiological studies based on worm expulsion and whole genome sequencing (WGS) is an obvious step in seeking to understand how transmission is sustained. WGS of expelled worms serves to reveal genetic diversity, and associated parasite population structure and relatedness. This information can be used to infer recent demographic changes ^20^, who infects whom by relatedness analyses, ^21^ the efficacy of interventions on diversity, and the associated impact of repeated treatment on single drug use ^22–24^. While a number of recent studies have looked at the population genetics of *A. lumbricoides* in endemic regions ^25–29^, the majority of these have employed a limited number of molecular markers with a focus on the relatedness of *A lumbricoides* to parasites in domestic animals such as pigs (e.g. *Ascaris suum*) ^27^.

In the context of infection control, the presence of multiple parasite genetic clusters within the host community and the relatedness between them may help identify transmission foci. Multiple sites of transmission, such as a local school serving many villages, or indeed specific households, can in principal be identified by molecular epidemiological studies based on WGS. While this approach has been adopted at a global geographic scale ^30,31^, to date, it has not been employed within a single community linked to longitudinal epidemiological studies of infection and control.

To elucidate the genomic impact of long-term MDA pressure on endemic communities, and to gain insights into the genomic diversity and gene flow amongst A. *lumbricoides* populations in an endemic community, we performed WGS on parasites collected during a worm expulsion survey in November 2022. We collected these worms as part of the Geshiyaro project ^11^, which is designed to measure the success of MDA and WaSH infrastructure improvements delivered with behaviour change communication sensitisation. The Geshiyaro project’s key objective is to interrupt STH transmission in the Wolayita region of Ethiopia. Using individual level epidemiological data alongside genomic data from chemotherapeutically expelled worms, we report the genetic structure of *A. lumbricoides* and patterns of transmission within and between households and villages. We also determined whether there are observable changes in STH diversity that could be attributed to repeated albendazole treatment. We also assessed the presence of variants which have been implicated in reduced albendazole efficacy ^32–34^.

To establish the effectiveness of the treatment programme, our analysis was disaggregated by age and drug compliance groupings (the fraction of the population who repeatedly take drug treatment) defined in the Geshiyaro control programme ^11,35^. We utilised household GPS data of participants to understand the presence of spatially explicit patterns of genetic associations between worms collected across the community ^36^. This study represents the largest WGS study of *A lumbricoides* and the first genome analysis collected within a large longitudinal study of infection levels linked to drug treatment.

## Materials and Methods

### Worm Collection

Worms were collected from 20 households within two Gotts (collection of houses within a community) in the Korke Doge (**Figure 1**) kebele (a collection of households clusters equivalent to a small village) after a routine deworming activity implemented by the Geshiyaro project. The study design and objectives have been described previously ^11,37^. Following treatment of 400 mg of albendazole delivered household-to-household, worms were collected from the stool of individuals for five days.

**Figure 1:**
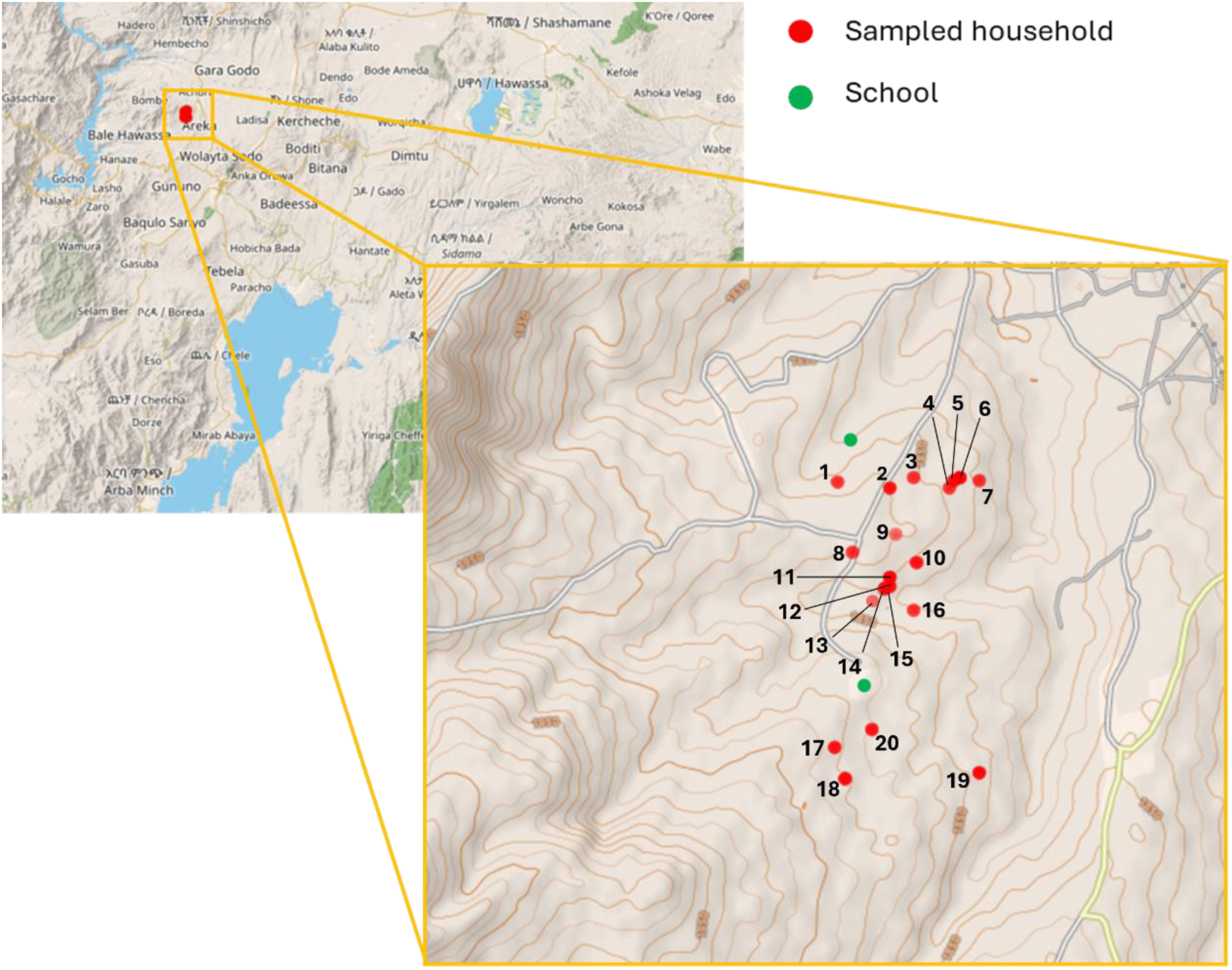
Map of sampled households across Korke Doge kebele in the Wolaita zone of Southern Ethiopia. Households sampled are denoted in red while school locations are highlighted in green. Household number included alongside location points.

### Epidemiological data collection

Annual parasitological mapping took place across 45 sentinel sites in 45 kebeles whereby stool samples were collected within the Geshiyaro cohort tracking infrastructure ^38,39^. All participants were enrolled to the project using electronic data capture fortified with biometric fingerprint technology for the registration and subsequent identification of participants. This generated an 11-digit participant number, which could be identified either biometrically, by scanning their ID card, or searching for their name.

By November 2022, Korke Doge sentinel site had received four rounds of community-wide (cMDA) since 2018. Albendazole was offered to members of the community aged one year old and above, with syrup distributed to infants aged one to four years old and pills to older individuals.

### Treatment compliance datasets

During cMDA, each participant was identified using their biometric fingerprint or ID card, which linked their 11-digit participant number to their treatment behaviour. Each participant was categorised as either ‘missed’ or ‘contacted’, with the latter further categorised as ‘accepted’ or ‘refused’ the offered drug, and subsequently ‘refused’ or ‘swallowed’ the accepted drug, as recorded by the HEW. The eligible population for albendazole was calculated as the population aged one year old and above, excluding all pregnant women. The term “fully compliant” defines individuals who have been categorised as ‘accepted and ‘swallowed’ albendazole at each round whilst the term “non-compliant” is refers to individuals who ‘refused’ at each round of treatment. The term “semi-compliant” defines individuals who ‘accepted’ albendazole at some rounds of MDA, but not all.

### DNA extraction and sequencing

DNA extraction was performed on snip sections of worm tissue from 54 adult worm samples. This was performed using the Qiagen MagAttract magnetic beads extraction methods. *A. lumbricoides* DNA samples were sequenced using Illumina NovaSeq (www.illumina.com) short-read paired-end sequencing, performed at Novogene Genomics (www.novogene.com). DNA was quantified by UV Spec and Picogreen. A 100 ng of DNA based on picogreen quantification was used as a template for NGS library preparation using the TruSeq Nano DNA Sample library prep kit. The mate-pair libraries were generated using the Nextera Mate Pair Library Prep Kit, following the gel-free method. The only modification to this protocol was that M-270 Streptavidin binding beads were used instead of M-280 beads.

### Variant discovery and annotation

Raw sequence reads from all 54 samples were examined using fastqc (v0.12.0) ^40^ to remove low-quality bases and adapter sequences. Trimmed sequences were aligned following the workflow of Easton et al. (2020) ^27^. Trimmed sequence reads were aligned using BWA mem (v0.7.18) ^41^. PCR Duplicates were marked using PicardTools (v3.3.0) MarkDuplicates ^42^. Variant calling was performed using GATK HaplotypeCaller (v4.0) in gVCF mode, retaining both variant and invariant sites. Variant sites with one single-nucleotide polymorphisms (SNPs) were separated from indels (insertion or deletion of bases in the genome) and mixed sites (variant sites having both SNPs and indels) using GATK SelectVariants. GATK VariantFiltration was used to filter both these independently. SNPs were retained when meeting the following criteria in the filtering process. VCFtools (0.1.16) ^43^ was used to exclude accessions with a high rate of variant site missingness (>5% of sites calling missing genotype). Subsequently sites were removed where >10% of accessions had a missing genotype. This filter process generated the primary VCF file for downstream analysis. To analyse nucleotide diversity (Pi) and fixation index (*F_ST_*), a second VCF file was produced. The depth of read coverage was calculated in 2 and 25kb windows along each chromosome using bedtools coverage (v2.19.1) ^44^.

### Population genomic structure and diversity

We removed all variants found to be in strong linkage disequilibrium. Principal component analysis was performed on the remaining 1,843,016 autosomal SNP using PLINK ^45^. Genomic structure was analysed using Admixture ^46^with *K* values (hypothetical ancestral populations) ranging from 1 to 20,

The 1,843,016 autosomal variants were used to construct a neighbour joining tree. An identity-by-state distance matrix was created with distances expressed as genomic proportions generated by PLINK. The ape bionj algorithm ^47^ was performed on the resulting matrix and visualised using ggtree ^48^. PIXY ^49^ was used to calculate the nucleotide diversity (π), fixation index (*F_ST_*) and absolute divergence (*d*_XY_), in 5 kb sliding windows with no-overlap across autosomes for each household and designated treatment compliance dataset.

### Spatial population genomics

To understand the extent of variation in genetic structure by location and identify spatial genetic discontinuities across the study community, several related methods were employed. Bayesian clustering with BAPS v5.3 ^50^ was used to test for the presence of multiple genetic populations under Hardy-Weinberg equilibrium, Principal components analysis (PCA) was employed to identify major trends in the genomic structuring through ordination. The fineSTRUCTURE method was used to assess genetic structuring through differences in shared co-ancestry ^51^. Spatial principal components analysis was employed to assess the genetic structuring by capturing patterns via spatial autocorrelation within the spatial genetic variability ^52,53^.

For detailed explanation of methods and workflows throughout analysis, please see Supplementary Text 1.

## Results

### Changes in the prevalence and intensity of infection over time

The baseline prevalence of *A. lumbricoides* in Korke Doge was 38.6% (95% CI: 30.7 - 46.6%) in Year 1 (2018), which decreased to 9.27% (95% CI: 4.63 - 13.9%) in Year 5 as illustrated in Figure 2a. Prior to the worm expulsion, the kebele received four rounds of community-wide MDA (cMDA) at a compliance rate (treatment swallowed) of 89.7%, 83.1%, 84.4% over three annual rounds, and a compliance rate of 87.7%, 73.9%, 60.0%, 68.9% over four subsequent biannual rounds.

**Figure 2:**
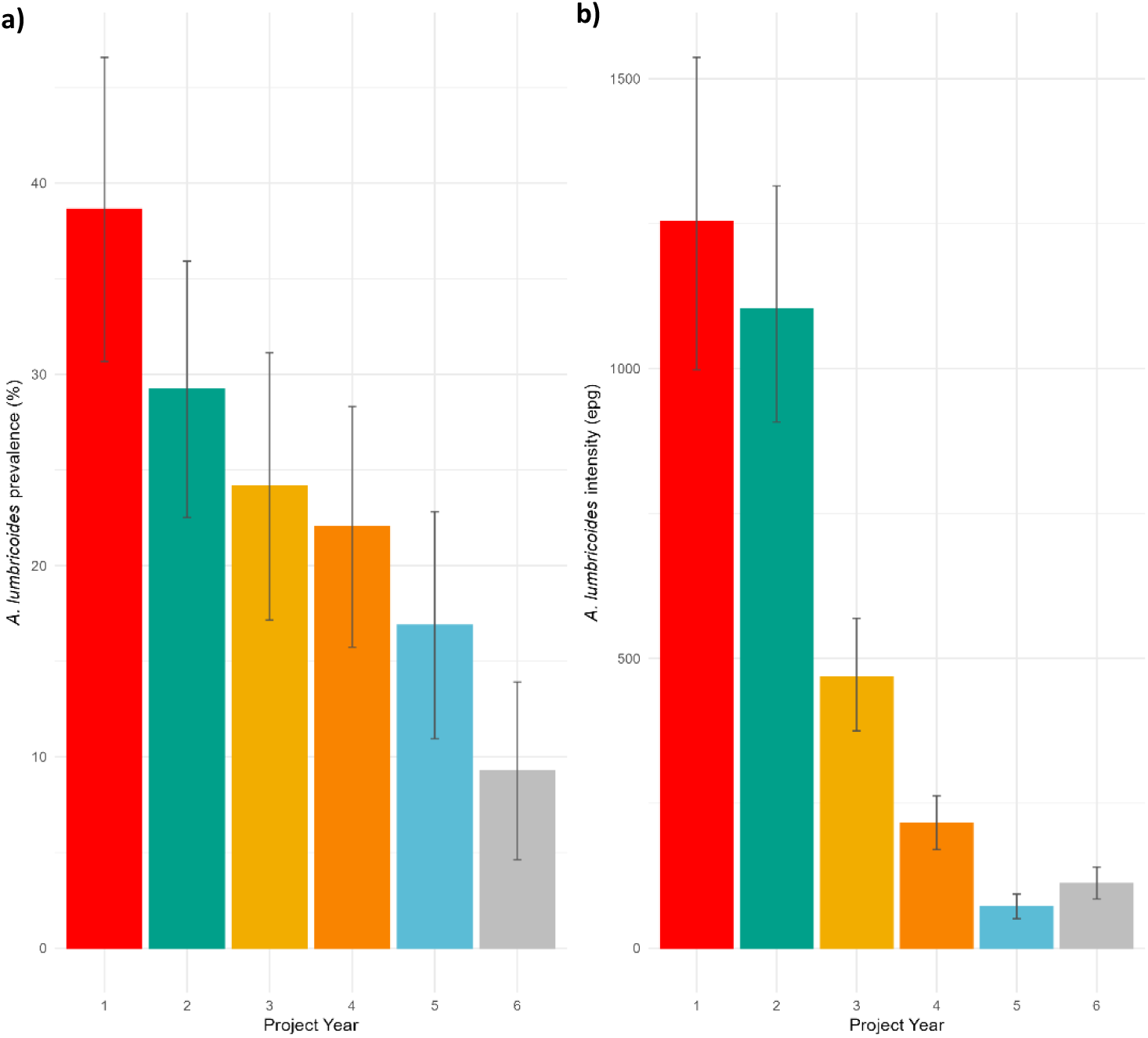
Change in **a** prevalence and b intensity (right hand axis) of *A. lumbricoides* in Korke Doge over five years of project implementation from 2018 to 2022. The target 2% prevalence of *A. lumbricoides* is shown as a grey dashed line. Error bars show the 95% confidence limits. All infections are classified as low intensity according to WHO guidelines.

The proportion of participants who remained infected between survey years decreased from 16.4% to 4.1%, as demonstrated in Table 1. The level of predisposition to no, light or heavy infection, as measured by Kendall’s tau ranking correlation statistic, in all the sentinel site cohorts in the Geshiyaro study increased from 0.18 (Year 1 to Year 2, p < 0.001) to 0.27 (Year 5 to Year 6, p < 0.001). For individuals in Korke Doge, predisposition also increased from a Kendall’s tau value of 0.19 (Year 1 to Year 2, p < 0.05) to 0.24 (Year 5 to Year 6, p < 0.01). Higher levels of predisposition to heavy infection were seen in individuals who were partially treated, compared with those who were always treated (Table 2).

**Table 1:**
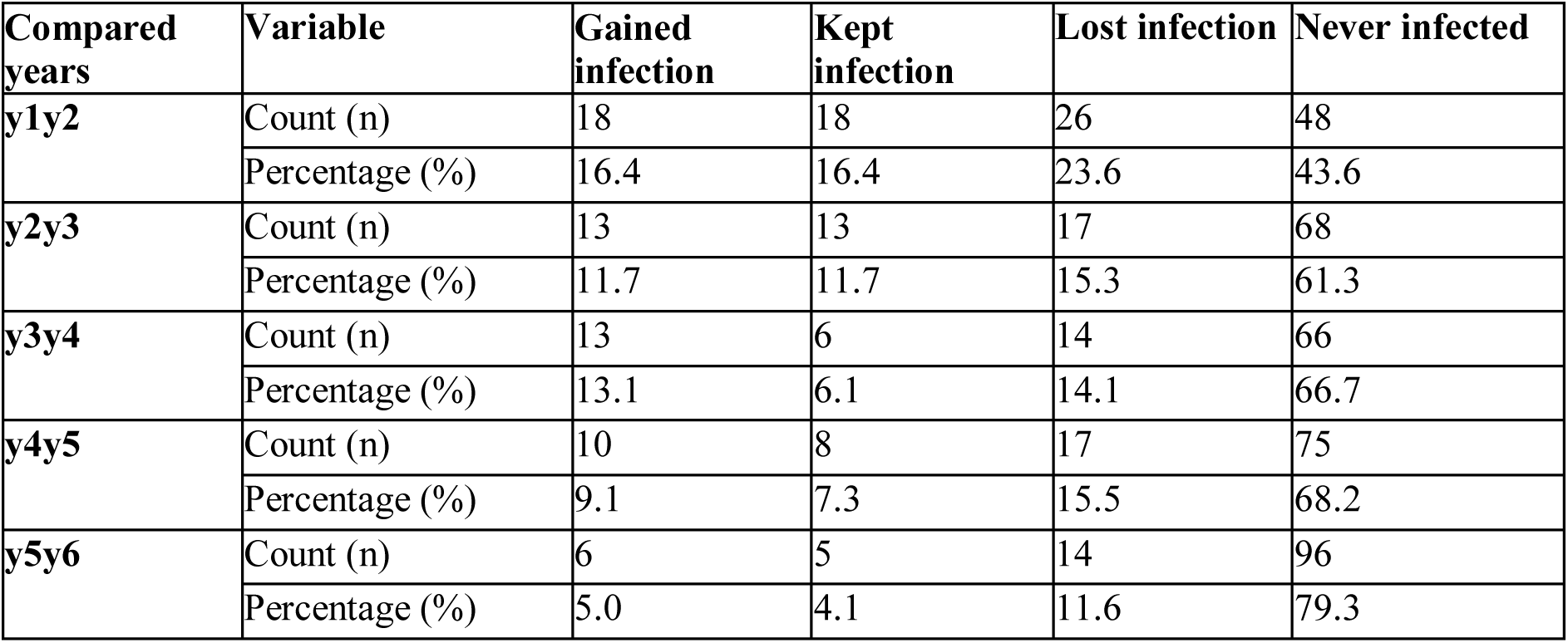
The year-on-year change in infection status in all surveyed individuals with the Korke Doge kebele.

**Table 2:** Predisposition to *A. lumbricoides* infection as measured by the correlation coefficient, Kendall’s tau. Correlation is measured on a scale from -1 to 1, where values close to 1 indicate a high level of predisposition to infection whereby those who were infected are more likely to be reinfected again. P values of significance are given as P < 0.001, p < 0.01, p < 0.05, or p > 0.05.

|  |  | y1y2 | y2y3 | y3y4 | y4y5 | y5y6 | y1y6 |
| --- | --- | --- | --- | --- | --- | --- | --- |
| **All**<br>**Geshiyaro**<br>**sites** | Overall | 0.180<br>P < 0.001 | 0.253<br>P < 0.001 | 0.215<br>P < 0.001 | 0.253<br>P < 0.001 | 0.271<br>P < 0.001 | 0.083<br>P > 0.05 |
|  | Always<br>treated | 0.065<br>P > 0.05 | 0.134<br>P > 0.05 | 0.170<br>P > 0.05 | - | - | - |
|  | Partially<br>treated | 0.187<br>P < 0.01 | 0.259<br>P < 0.01 | 0.178<br>P < 0.01 | 0.111<br>P < 0.01 | 0.178<br>P < 0.01 | 0.098<br>P > 0.05 |
| **Korke**<br>**Doge** | Overall | 0.194<br>P < 0.05 | 0.277<br>P < 0.001 | 0.117<br>P > 0.05 | 0.233<br>P < 0.01 | 0.242<br>P < 0.01 | 0.213<br>P < 0.05 |
|  | Always<br>treated | 0.098<br>P > 0.05 | 0.128<br>P > 0.05 | 0.118<br>P > 0.05 | - | - | - |
|  | Partially<br>treated | 0.177<br>P > 0.05 | 0.269<br>P < 0.01 | 0.105<br>P > 0.05 | 0.259<br>P < 0.05 | 0.218<br>P < 0.05 | 0.206<br>P > 0.05 |

### Worm burden

A total of 102 individuals were sampled across the community over five days following community-wide albendazole treatment. Of these 102 individuals, 54 individuals were found to expel at least one adult worm, with 222 worms expelled in total (Supplementary Table 1). Worm count per person showed a slight increase in Pre-SAC and SAC individuals, but this was not statistically significant (Supplementary Fig 1). Of the 54 individuals that expelled worms, 31 were from non-compliant individuals, 11 from semi-compliant and 8 from fully compliant (Supplementary Table 1).

### Whole-genome sequencing of individual worms

Alignment of sequence reads to the reference *A. lumbricoides* genome revealed high-sample mapping rates (median 87.99% of reads mapped with the mapping quality MQ>40 (Accession SRR31675196 - SRR31757547; Supplementary Table 2). Reads mapped to the majority of the reference genome (median of 94.23% of bases covered across all samples), although the median read depth was highly variable across the total sample of worms (1.33-50X). Across all worms, a single worm was taken per individual for a final analysis dataset of 54. The final variant dataset comprising of 3,692,001 single-nucleotide polymorphisms (SNPs) and 280,447 indels (including mixed SNP/indel variant sites) from 54 adult worms.

### Local parasite population genetic structure

The population structure of the 54 worm sample is depicted by a variety of analyses in Figure 3. We generated a maximum likelihood phylogeny (Figure 3a) and a principal component analysis (PCA) using a subset of 1,000,000 unlinked autosomal variants (Figure 3b). PCA indicated three genetic clusters existed within the infected households in the sampled community (Figure 3b and c). ADMIXTURE analysis consistently differentiated parasites within Korke Doge households with cross-validation analysis (Supplementary Fig 3) of? the most likely number of subpopulations (*K*) and confirmed this for a range of *K* values. Specific differences in inbreeding coefficients are observed across households 1, 3, 12, 13, 14, 15, 17, 19 and 20 in Figure 4 indicating reduced geneflow between these households and others within community. Pairwise measures of fixation revealed low levels of diversity across all households (Supplementary Table 3). Across the community we establish summary values of HO = 0.212, HE = 0.405, p = 0.395 and FIS = -0.121 for the full data set (Supplementary Fig 4).We characterized the population structure at different spatial scales using a subset of 1,000,000 autosomal variants. These were filtered to remove variants in strong linkage disequilibrium. Principal component analysis revealed that 34% of total variance was explained through the first two components. Phylogenetic analysis revealed no real discernible structure within parasites across Korke Doge apart from the three distinct genetic groupings of the parasite shown in Fig. 5a & b. Genome-wide estimates of median nucleotide diversity within parasite population revealed low levels of differentiation (π = 0.02, *d*_XY_ = 0.09; Supplementary figure 4). Parasite populations within households showed approximately equivalent levels of nucleotide diversity (π = 0.01, *d*_XY_ = 0.05) (Supplementary Fig 4).

**Figure 3:**
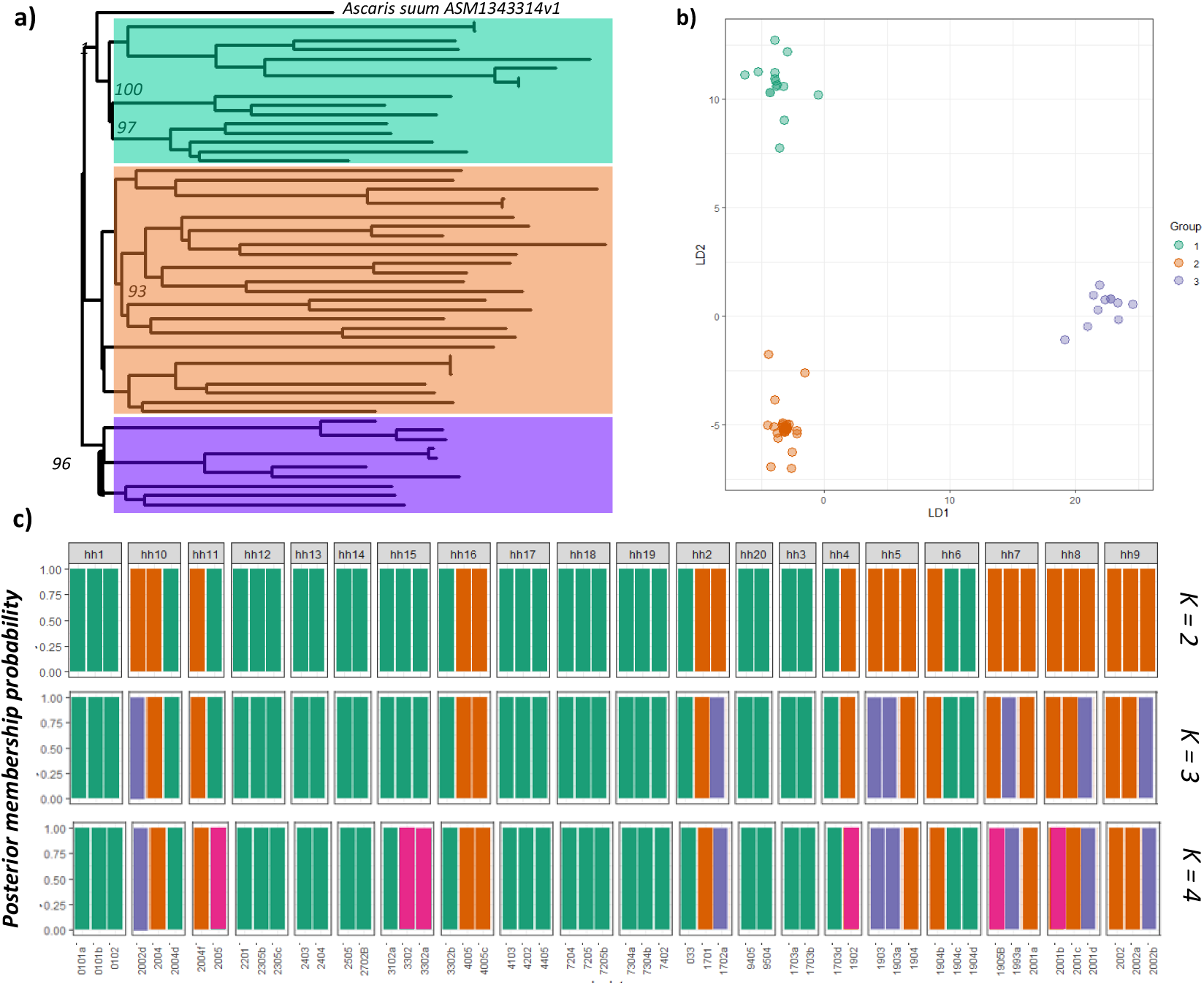
*Ascaris lumbricoides* population structure. Principal component analysis (PCA) of genetic differentiation within and between the 54 *A. lumbricoides* produced sampled for WGS **a** Neighbour-joining phylogeny showing relatedness between samples, branches are coloured according to principal component clustering. **b** Principal components 1 and 2 accounting for approximately 34% of total variance. **c** cluster assignment illustrating the cluster assignment per individuals per household. **c** ADMIXTURE plots illustrating the population structure, assuming 2-4 populations are present (*K*), using 10-fold cross-validation and standard error estimation with 500 bootstraps. *Y-axis* values show admixture proportions for different values of *K* (*K =* 2-4), each colour indicates a different population.

**Figure 4:**
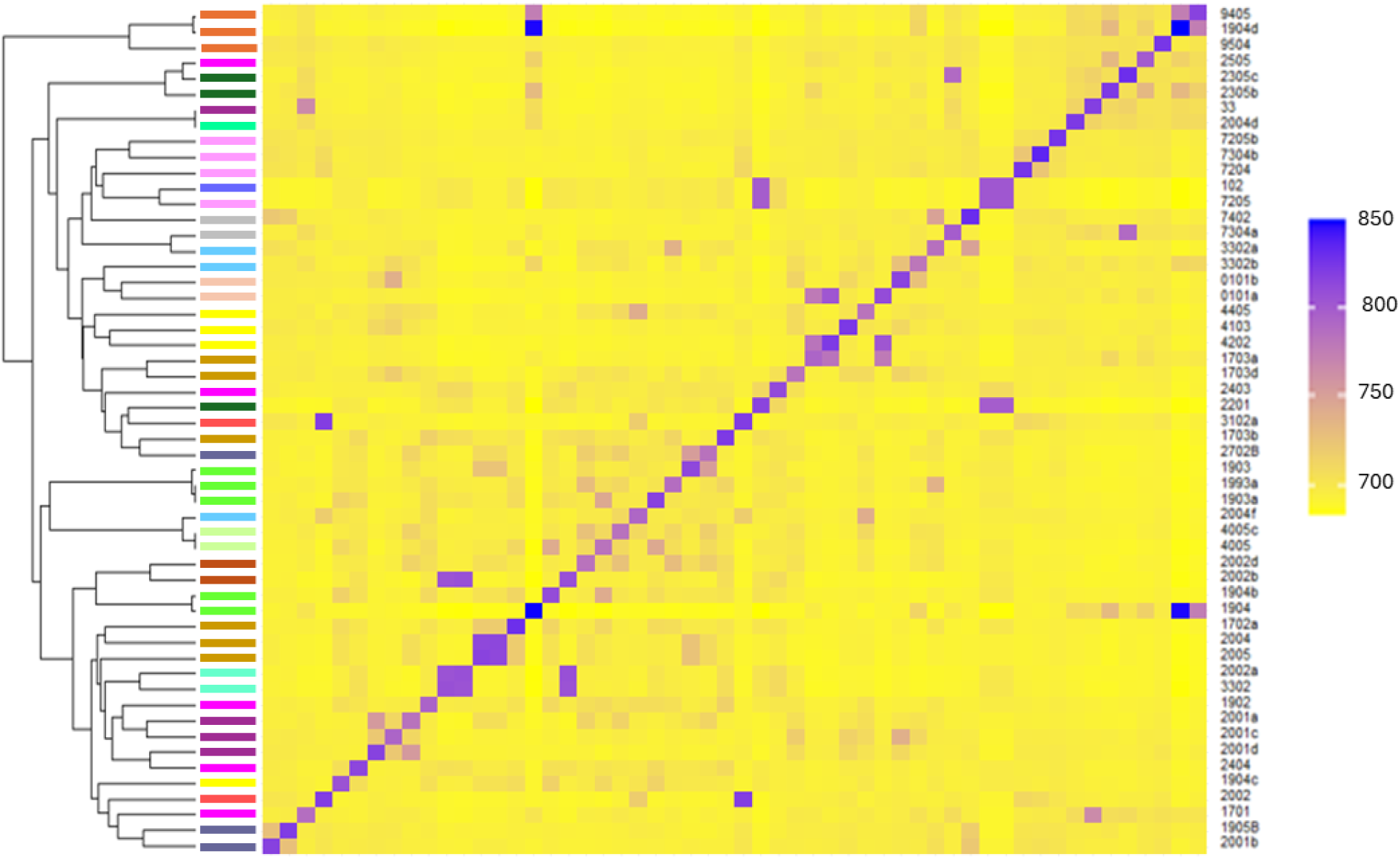
Co-ancestry heatmap. Co-ancestry heatmap generated through fineSTRUCTURE analysis. Samples grouped along the heatmaps diagonal have common shared co-ancestry histories and pairwise comparisons outside the diagonal indicate level of co-ancestry between individuals. Lighter yellow represents lower shared co-ancestry and shades of purple indicate progressively higher shared co-ancestry. Colours of corresponding bands alongside the dendrogram indicate household membership in which each worm was sampled.

**Figure 5:**
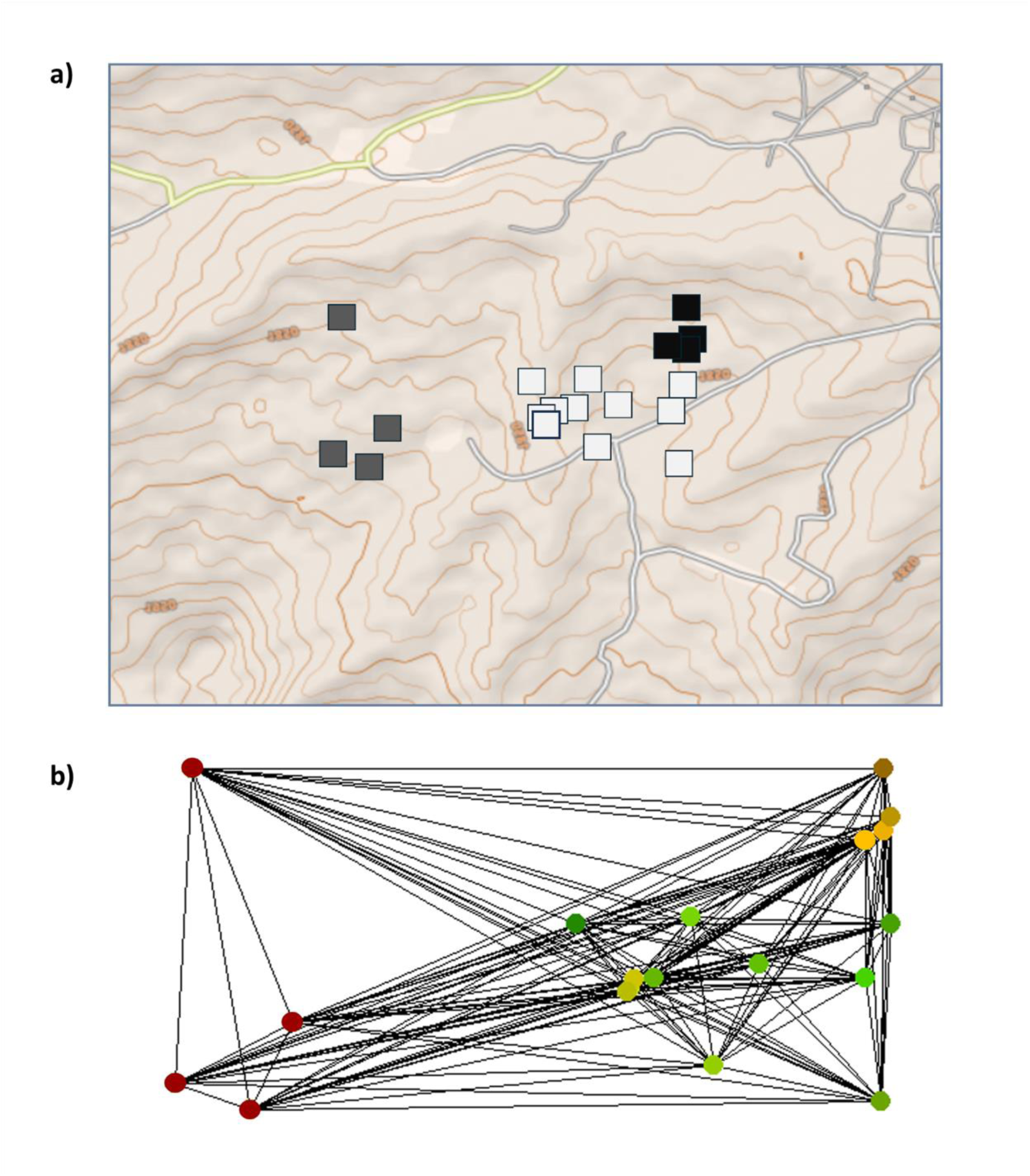
**a** Represents the first global axis scores from full landscape sPCA (*n* = 54). Each georeferenced household is depicted with a square, where dark grey colour represents the major spatial genetic clusters of sampled households. **b** network used to define spatial weightings for sPCA analysis, colours indicate the spatial genetic cluster assignments.

### *Ascaris lumbricoides* population size within the community following multiple rounds of community-wide MDA

We estimated an effective parasite population size of 201.3 (95% CI 197.1 – 205.5) across the entire community. The dataset was further disaggregated by an individuals’ compliance to the annual treatment programs, where compliance is taken to mean the swallowing of the offered treatment. The effective population of worms within different compliance groups were as follows; fully compliant 29.9 (95% CI 21.3 – 38.2), semi-compliant 98.5 (95% CI 80.7 – 106.3) and non-compliant 121.3 (95% CI 115.8 – 136.8). Additionally, the community was stratified into defined age groups; namely Pre-SAC (2-4 years), SAC (5-14 years), Adolescents (15-20 years), young adults (20-35) and Adults (36+). The effective population size of parasites in these age groups were estimated to be; Pre-SAC 88.2 (95% CI 54.8 – 121.6), SAC 135.6 (95% CI 119.5 – 151.7), Adolescents 45.2 (95% CI 36.1 – 51.3) and Adult 57.5 (95% CI 40.9 – 64.1) (Supplementary Fig 4). In summary, most worms are predicted to be in SAC, and within that group, the majority are in those non-compliant to treatment (Supplementary Table 1).

Genome-wide allele frequency patterns allow the determination of recent reductions in population size and have been demonstrated as a reliable metric for establishing demographic size change in helminth populations^24^. Analysis of one-dimensional site frequency spectra (1D-SFS) across the Korke Doge sample does not indicate an abundance of rare (singleton and doubleton) alleles which suggests the parasite population effective population size is small and declining (Supplementary Fig 5a). Median genome-wide Tajima’s D estimates were positive across all compliance groups which suggests a recent population constriction most likely influenced by the past six years of MDA (Supplementary Fig 5b). Demographic size change indicated that the effective population size of parasite populations over time, revealing current populations are a fraction of the size of the historical populations from which they were derived (Supplementary Fig 5c).

### Community-wide spatial population genomics

We identified signals of population differentiation using methods that aim to identify fine-scale genetic structuring. Co-ancestry heatmaps between individual worms (Figure 4) highlighted reduced geneflow between worms from compared from within households compared to between. Pairwise comparisons among these households reveal a pattern of co-ancestry with increased gene flow between closely located households and decreased geneflow with worms in the more distant households.

Plotting values from the first axis of the sPCA analysis indicated a break in connectivity across these spatially distinct clusters. In Figure 5a. Grey scale squares indicate households with spatially discriminate clusters of worms. Figure 5b shows the results of spatial PCA analysis, these are a composite measure of both genetic diversity (variance) and spatial structure (autocorrelation through Moran’s I). λ_1_ scores from spatial PCA also show a distinct spatial structure across households within sampled community (Figure 6). Colour assigned spatially distinct clusters indicate that neighbouring households demonstrate reduced variance between them, with increasing variance occurring within households at greater distance. Results have been plotted across the Delauney triangulation connection network used to calculate spatial weightings with each node within the network representing a sampled household. Plotting of eigenvalues resulting from sPCA revealed a pattern of local scale structuring (Supplementary Fig 5). Plotting of interpolating lagged principal scores show a map of genetic clines between the sampled households in the community (Supplementary Fig 6). Spatial clustering was confirmed through spatial autocorrelation output from Moran’s I (Supplementary Fig 7).

**Figure 6:**
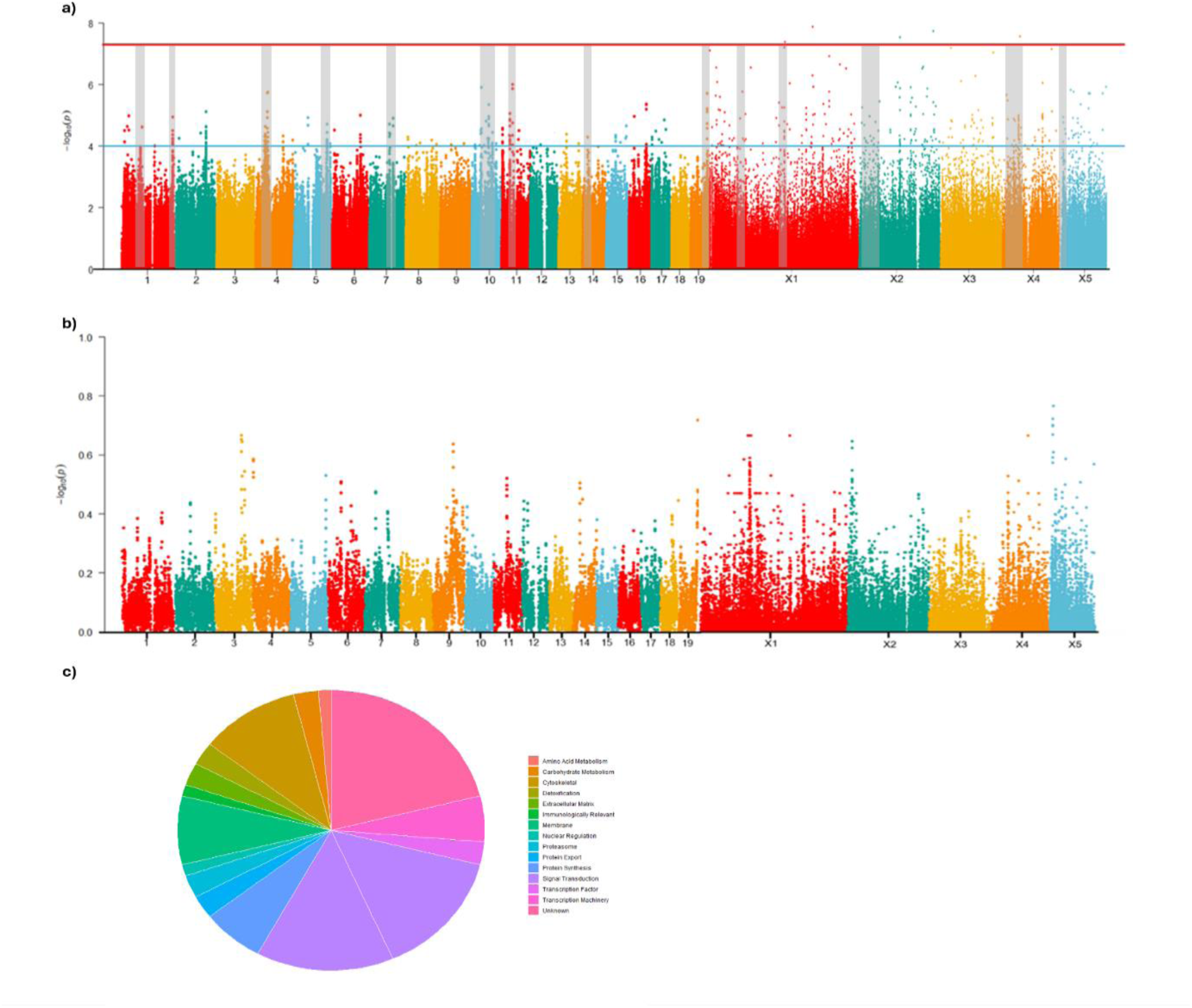
**a** Genome-wide integrated haplotype scores (iHS) within the Korke Doge population of expelled worms (54). Regions that show increased iHS scores in conjunction with elevated *F_ST_* highlighted within the grey filled boxes. **b** Fixation index *F_ST_* between fully compliant individuals which have accepted the drug each round of treatment and non-compliant individuals which have accepted treatment at only the latest round of treatment) within Korke Doge. **c** Pie chart of gene functions for genes in candidate regions of positive selection.

### Evidence of positive selection in a region that has experienced long-term MDA

A number of genome-wide methods were used to identify regions of the genome undergoing positive selection in worms from Korke Doge following multiple rounds of MDA. Multiple regions were identified by the integrated haplotype score (iHS) test as being under strong positive selection in Korke Doge (Figure 6a & b). Overall, we identified 69 non-redundant regions with extreme iHS scores (Supplementary Table 3). From these 69 regions we found 14 regions that indicated both elevated iHS and *F_ST_* values. We also determine whether any of these selected regions intersected with regions of reduced genetic diversity (Supplementary Fig 8). These regions were defined as encompassing genes which contained at least one variant in one or more worm samples ^54^. Candidate regions of extreme iHS scores spanned 91 protein coding loci with known functionality with a further 38 classified as hypothetical proteins (Supplementary **Table 3**).

One region that displayed strong signals of selection was located within chromosome 1. Across chromosome 1 there was three candidate regions (CR1 1.13 – 1.15Mb, CR2 2.58 – 2.6 Mb, CR3 7.27 – 729 Mb and CR4 17.52 – 17.55 Mb) which demonstrated increased levels of localised selection (Figure 7). Across these regions CR3 and CR4 demonstrated significantly elevated iHS values, increased *F_ST_* and decreased relative nucleotide diversity. Within the region on chromosome 1 we identified a single locus containing the beta-tubulin gene (AgB04_g300) which shows three non-synonymous variants at low to mid frequencies (0.11-0.42, Supplementary Table 3) at amino acid positions 7, 9 and 67. Within these regions we identified a single locus containing beta-tubulin gene. Additional beta-tubulin genes were within localised regions of selection in Chr X3 (7.87 – 7.877 Mb). However no non-synonymous mutations were identified in this region (Supplementary Table 3). The beta-tubulin families are implicated in reduced susceptibility of *A. lumbricoides* to anti-helminthics within the benzimidazole family ^32,55,56^. There was no evidence for specific mutations at three codon positions ^57,58^ previously linked to resistance against albendazole in *A. lumbricoides.* ^57,58^

Further evidence of positive selection was found across chromosomes 10 (CR20 1.14 – 1.16 Mb, CR21 3.65 – 3.67 Mb, CR22 3.89 – 3.91 Mb, CR23 4.81 – 4.83 Mb, CR24 4.95 – 4.99 Mb, CR25 6.22 – 6.24 Mb 7.47 – 7.72 Mb) which encompassed 19 genes. A variety of other genes in candidate regions encode non-synonymous mutations including a microtubule associated protein (AgR001_g156), dynactin, annexin, and transcription factors among others (Supplementary Table 3). A potassium voltage-gated ion channel protein was located within candidate region 27 (7.59 – 7.63 Mb).

## Discussion

To sustainably interrupt transmission of STH there needs to be a greater understanding of how these parasites sustain circulation in low prevalence communities. We performed WGS of individual *A. lumbricoides* parasites collected within a cohort-tracked, epidemiologically relevant framework across a single community. The first time that such analysis has taken place within an expanded, community-wide treatment programme. The study of one village and a relatively small sample of worms where WGS was possible (54 worms from an estimated total population of over 200 worms (∼25%) revealed several insights into transmission. First, genome data suggests that *A. lumbricoides* displays fine-scale population structure within a village with endemic infection (Figure 3 & 6). Secondly, multiple infective foci are present within the study community and finally individuals within the village are not being infected by a relatively homogenous randomly mixed population of worms.

The data suggests that transmission primarily occurs at the household level rather than single source community-wide infection. Genetic relatedness is strongest between closely located households rather than those more widely spatially distributed. The focal infective nature of *A. lumbricoides* infection within endemic regions and communities has been suggested from parasitological studies of infection patterns in Nepal ^30^, Venezuela ^59^, and Guatemala ^60^. Principal components analysis across our spatial network indicate a strong signal for local-scale structuring, inferring that genetic clusters are spatially explicit. The increase in homozygosity within these spatially explicit groups of worms suggests that directional selection will exhibit greater efficiency in driving an increase in potentially advantageous alleles across populations, such as drug-resistance genes within a population of frequently treated worms ^61^. As prevalence is driven down by numerous rounds of treatment understanding these patterns of spatially clustering would permit a more targeted drug administration strategy once clusters are identified. genomic clustering was not correlated with human host-specific factors, such as age, sex or treatment compliance. These variables certainly influence infection levels but not to the degree of genetic relatedness between worms in this community. Further investigation within these spatially distinct clusters into WaSH and compliance status is required to unravel the factors influencing the continued presence of specific transmission foci.

Parasite genomic data indicates a shrinking population of *A. lumbricoides* across Korke Doge. This reflects the cMDA treatment history of the region. Biometric tracking of participants has allowed the programme to collect individual-level data on cMDA compliance and infection status. While worm expulsions indicated that partially and non-compliant individuals harbour a much larger fraction of the total parasite population (Supplementary Table S1) overall, these worms are part of a shrinking population due to the years of MDA treatment in the community. The individuals who fall into this non-treated category are therefore the main infection reservoir in the community following cessation of expanded treatment programme.

In recent years evidence of reduced efficacy of albendazole has arisen in some settings with endemic infection and regular MDA ^62,63^. The nature of mutations that would be classified as rare resistance-associated variants at a single locus to be swept to fixation within a population would be classified as a hard selective sweep. These would leave a strong selective signature in the genome around the locus which can be detected by several methods such as the *F_ST_* and haplotype tests (iHS) that have been employed in this study. Other scenarios, such as the presence of a resistance allele across a range of genetic backgrounds, which subsequently would rise to gradually elevated frequencies (soft sweep) would be harder to detect and may have been missed in this study. This type of soft sweep would be more likely to be detected across a multi-community study across a larger geographic region. If variants associated with a range of genetic backgrounds, this would be demonstrated through a continued genetic variation across a longitudinal study, which has yet to be performed across soil-transmitted helminth species. A small number of studies have investigated the presence of resistance-associated mutations in *A. lumbricoides* collected as part of control programs, finding them occurring at low frequency (0.05%) ^64^. Alternatively, geographically disparate studies in Kenya, Haiti and Panama indicated a high proportion of the F167Y (position 200) resistance associated mutation^33^. Worms collected as part of this study did not display any specific substitutions related to albendazole resistance in line with global screens for the variant ^65^. The study community showed a high proportion of infected individuals being designated as “Lost infection” in subsequent annual survey following accepted treatment (Table 1). Therefore, albendazole still shows good efficacy across this community.

The main conclusions of the study are twofold. First, the study of one *A. lumbricoides* population showed fine-scale spatial division across households and sets of households that have received repeated cMDA. Transmission seems to be very household focused and genetic distance between worms rises as the spatial distance between households increases. Second, at present, despite long term repeated treatment, there appears to be no indication of resistance to albendazole treatment in the parasite population. However, we did identify several genomic regions with strong signals of selection that could indicate gradual adaptation to long-term albendazole exposure or other selective pressures. Further, much larger scale genomic surveillance across endemic regions, particularly alongside control programmes utilising tracked cohorts is clearly desirable.

In low prevalence communities, the key policy question at present is how best to focus control efforts. Improved WaSH is essential in terms of its wide impact on many infections. Continued prophylactic cMDA of large populations, when only a few are infected, is less clear cut. MDA alone at very high coverage repeatedly for many years can stop STH transmission, but the expense and the continued availability of drug donations in resource poor settings is less certain. Moreover, this repeated use of single drug treatment on mass-scale without understanding methods to monitor specific loci involved in drug resistance, will put future programs at risk ^66^, particularly in poorly delivered, expanded programmes. Targeting treatment to the few who remain infected, many of whom are predisposed to infection, is one possibility as this study suggest that the household is the key basic unit of transmission for *A lumbricoides*. However, the use of mass diagnosis by parasitological stool examination or other more sensitive methods ^67^ is also expensive, certainly if repeated for all MDA rounds. Predisposition and the identification of household hotspots of transmission could significantly reduce the workload and associated expenses since once identified in low prevalence settings treatment would in the future just focus on these households. Molecular epidemiological approaches are widely used in the study of the epidemiology and control of viral ^68^ and bacterial ^69^ pathogens but are very rare in the study of the neglected tropical diseases. The costs of sequencing large genomes continue to reduce and hence larger-scale studies of ‘whom infects whom’ and routine genetic surveillance will become more feasible. It is hoped that molecular epidemiological approaches within a framework performed here will be employed more widely to further understanding of transmission in low prevalence settings.

## Supporting information

Supplementary Figures Doc

## Acknowledgements

We thank the technicians, local health workers from the woreda health division in Wolaita. We would also like to thank the technicians involved in the collection of samples in Korke Doge kebele: Selamawit Ginjo, Silas Dea, Moges Mekonin, Selamu Dana, Simegn Mengistu, Abraham Anjulo, Michael Tumato and Misgana Zekarias. We would also like to thank the community leaders and members of the Korke Doge kebele who were recruited into the study and their cooperation with everyone involved.

## Notes

### Competing Interest Statement

The authors have declared no competing interest.

### Summary of Updates

Adjusted text formatting, author affiliations updated

