## Supplementary Figures Doc for "Molecular epidemiology of *Ascaris lumbricoides* following multiple rounds of community-wide treatment: who infects whom?"

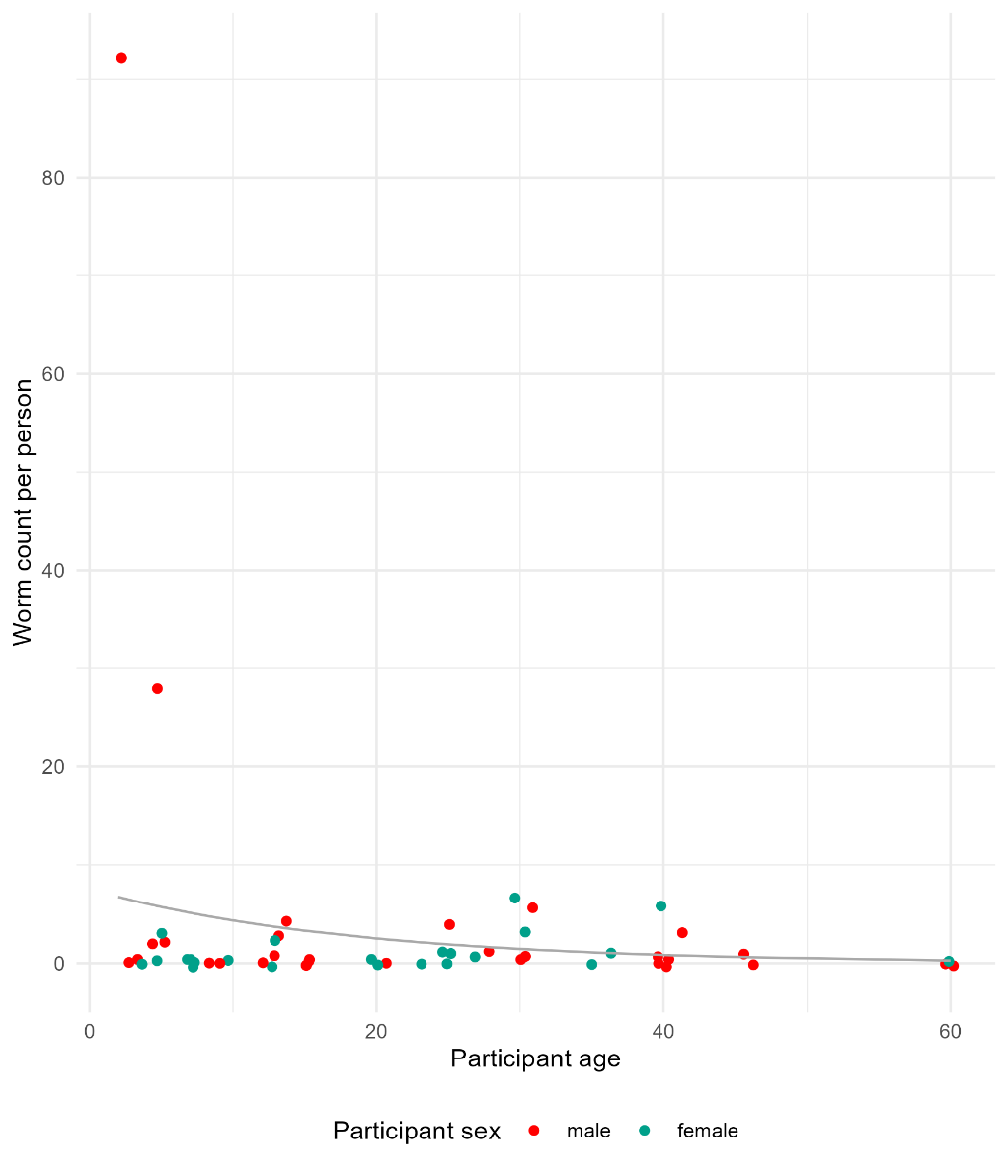

**Supplementary Figure 1:** Counts of worms expelled through chemo expulsion. Increasing age is displayed across the *x-*axis and worm count per individual is displayed in the *y*-axis. The worm count is the sum of total number of worms collected across five days following albendazole ingestion

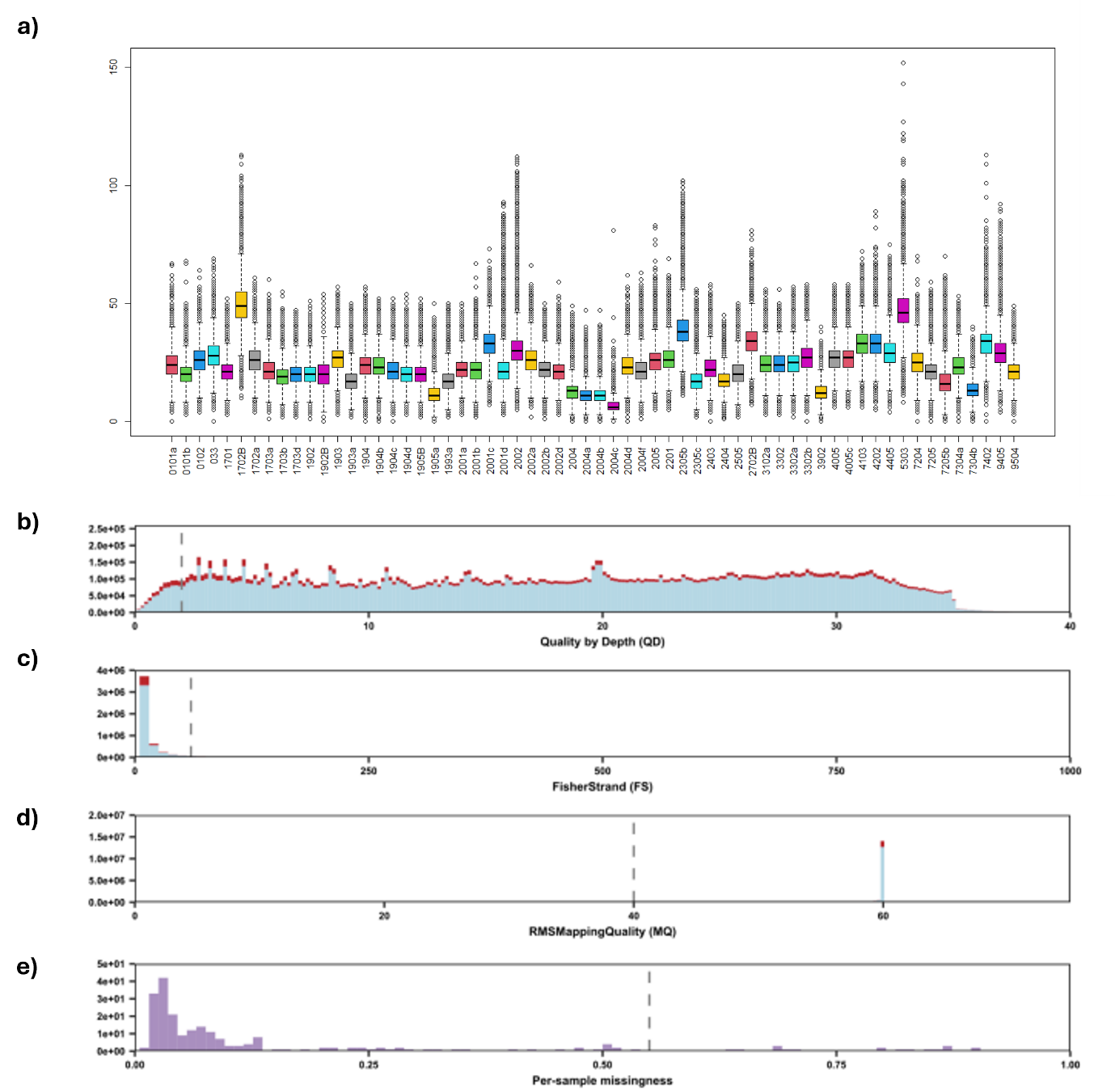

**Supplementary Figure 2: Variant quality control.** Plot a shows the mean depth of coverage per sample. Plots b and c show the frequency distribution of mapping values throughout the 3,692,001 variants. Plots d and e show the frequency distribution of per-sample missingness (samples with a high rate of per-site variant missingness) and per-site (sites with a high proportion of variant missingness) missingness after filtering using thresholds. Vertical dashed lines show the thresholds applied for the removal of sites.

| K | log(ml) |
| --- | --- |
| 1 | -15005054.776 |
| 2 | -1497772.129 |
| 3 | -14800534.39 |
| 4 | -1492882.662 |
| 5 | -1506327.448 |
| 6 | -1508411.942 |
| 7 | -1513480.551 |
| 8 | -1518480.551 |
| 9 | -1519967.927 |
| 10 | -1521112.44 |
| 11 | -1533352.44 |
| 12 | -1551002.44 |
| 13 | -1553228.58 |
| 14 | -1554773.25 |
| 15 | -1554997.114 |
| 16 | -1560001.287 |
| 17 | -1565551.88 |
| 18 | -15633147.11 |
| 19 | -1577821.02 |
| 20 | -1580357.13 |

**Supplementary Figure 3: Log maximum likelihood values** calculated by BAPS for 1-10 putative populations (K) across the full filtered dataset (*n* = 54)

| **Data set** | ***n*** | ***H*_O_ (±StdErr)** | ***H*_E_ (±StdErr)** | **Pi (±StdErr)** | ***F*_IS_ (±StdErr)** | ***N*_e_ (95% C.I.)** |
| --- | --- | --- | --- | --- | --- | --- |
| Total dataset | 54 | 0.212  (0.001) | 0.405  (0.001) | 0.395  (0.001) | -0.121  (-0.077) | 81.6 (67.1–95.1) |
| **Age groups** | | | | | | |
| Pre-SAC | 9 | 0.207 (0.023) | 0.251 (0.026) | 0.352 (0.026) | -0.168 (-0.048) | 88.2 (54.8 – 121.6) |
| SAC | 18 | 0.200 (0.022) | 0.260 (0.026) | 0.264 (0.026) | 0.221 (0.073) | 135.6 (119.5 –151.7) |
| Adolescents | 14 | 0.204 (0.024) | 0.245 (0.027) | 0.246 (0.027) | -0.159 (-0.043) | 45.2 (36.1 – 51.3) |
| Adult | 13 | 0.188  (0.029) | 0.208  (0.031) | 0.338  (0.029) | -0.099  (-0.088) | 57.35  (40.9 – 64.1) |
| **Compliance** | | | | | | |
| Fully Compliant | 8 | 0.103  (0.001) | 0.121  (0.005) | 0.127  (0.003) | -0.157  (-0.055) | 29.9 (21.3 – 38.2) |
| Semi-compliant | 11 | 0.199  (0.005) | 0.287  (0.005) | 0.329  (0.002) | 0.210  (0.029) | 98.5 (80.7 – 106.3) |
| Non-compliant | 31 | 0.279  (0.055) | 0.375  (0.039) | 0.377  (0.009) | 0.255  (0.087) | 121.3 (115.8 – 136.8) |

**Supplementary Figure 4: Population genetic indices across dataset.** This table describes the population genetic indices across the sampled community. It is disaggregated via age grouping and drug compliance throughout Ho, He, Pi, Fis and Ne for each group.

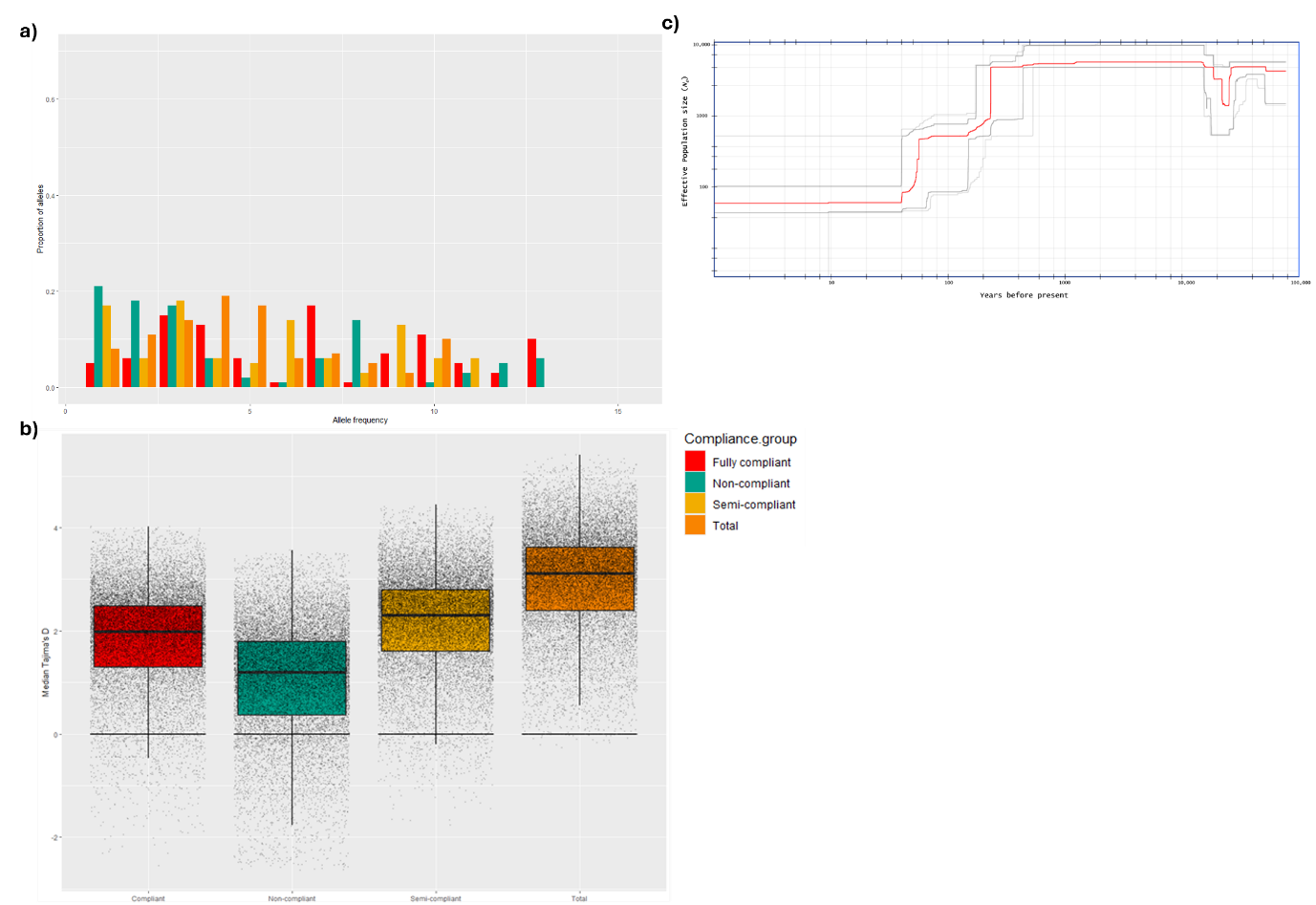

**Supplementary Figure 5: a** One-dimensional site frequency spectra for each parasite population according to drug compliance assignment. The *x*-axis represents the derived allele frequency and the *y*-axis represents the proportion of sites at each allele frequency. **b** Median Tajima’s D values calculated in 5 kb windows across each autosome for each compliance group. For all boxplots the central line indicates the median the top and bottom edges of the box indicate the 25^th^ and 75^th^ percentiles, respectively, the maximum whisker lengths are specified as 1.5 times the interquartile range. **c** Demographic history change within the Korke Doge population

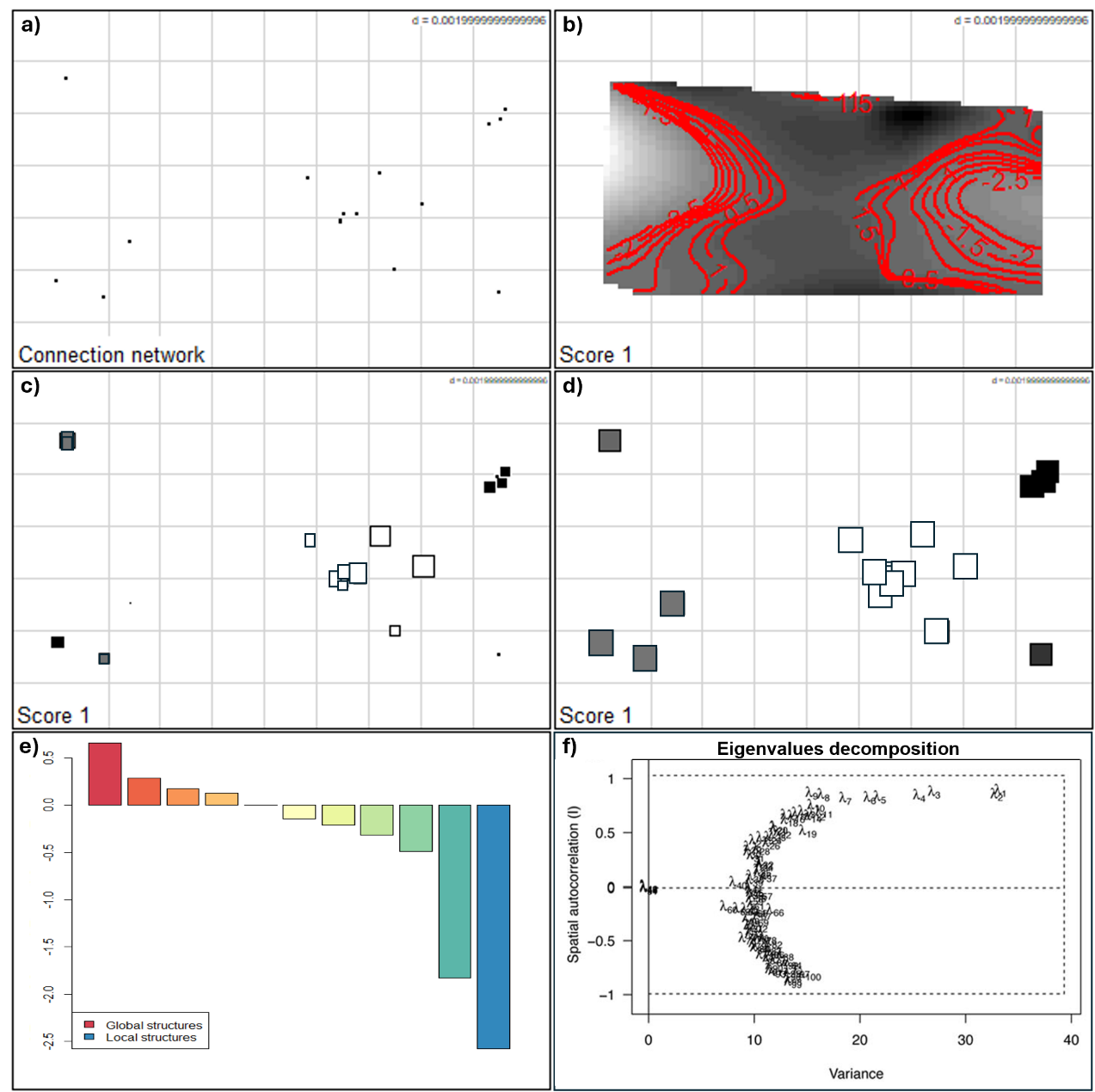

**Supplementary Figure 5: Spatial Principal Component analysis**: **a.** represents the points used in the connection network that was used to define spatial weightings. **b-d** plots representing the first eigenvector scores in space, **b** showing a contour plot representing the plotting of $li local scores, the closer the contour lines the greater level of genetic differentiation is in space. **c** plotting the local scores in greyscale, large black squares are well differentiated from large white squares, small squares are less well differentiated from each other. **d** This plot is a variant on grey levels. All three plots taken in the round indicate that three genetic clusters exist in three genetic clusters. **e** plot represented the local and global score eigenvectors. **f** Represents eigenvalues of sPCA denoted λi with i = 1, . . . , r, where λ1 is the highest positive eigenvalue, and λr is the highest negative eigenvalue according to the variance and Morans’s *I* components

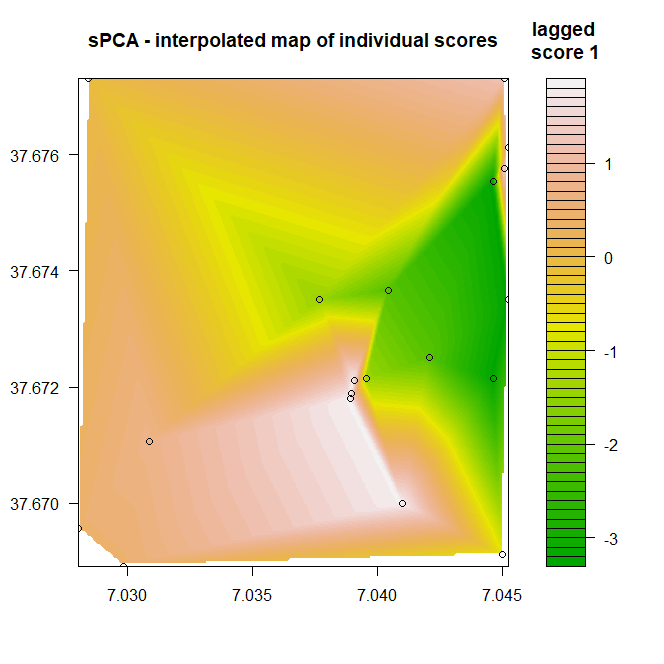

**Supplementary Figure 6:** Interpolated map; map of principal components onto geographic space. To achieve better resolution the lagged scores have been plotted on specific interpolated coordinates. Each circle represents a sampled household

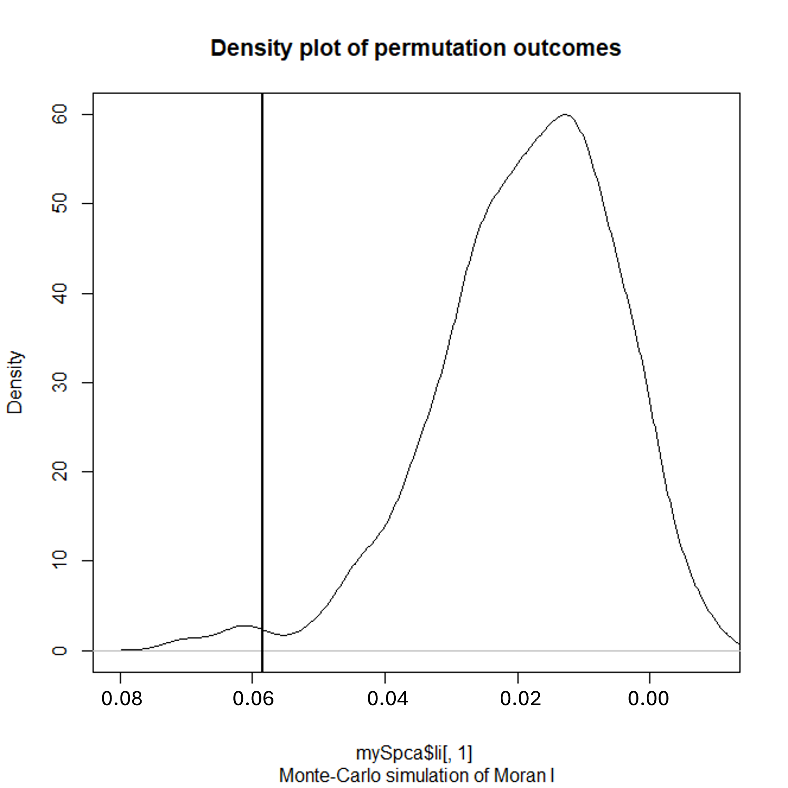

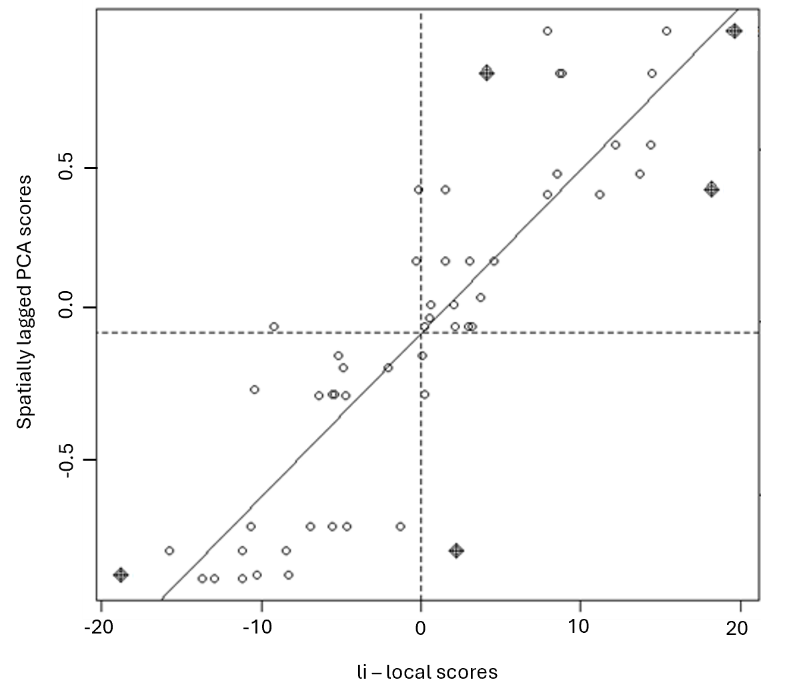

**b)**

**a)**

**Supplementary Figure 7: a.** Negative autocorrelation outcome plot between spatial variable and its lag factor. **b.** Density plot of Moran’s I posterior scores

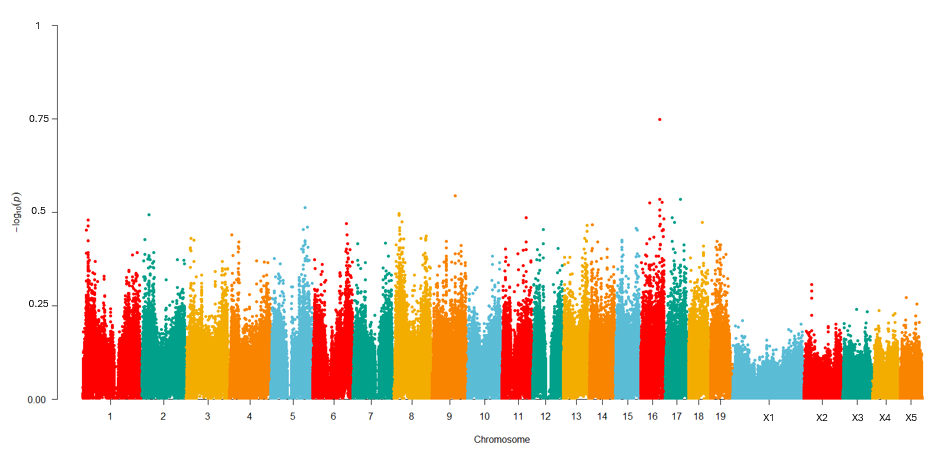

**Supplementary Figure 8:** Genome-wide plot of genetic diversity (Pi) from all individuals within the dataset
