## Supplementary Text 1 for "Molecular epidemiology of *Ascaris lumbricoides* following multiple rounds of community-wide treatment: who infects whom?"

**Epidemiological data collection**

Annual parasitological mapping took place across 45 sentinel sites in 45 kebeles whereby stool samples were collected, and participants answered a WaSH infrastructure access questionnaire. A demographically balanced sample of 150 participants were selected per site, consisting of 15 individuals of each sex in each age group: preSAC (aged 1-4 years), SAC (aged 5-14 years), adolescents (aged 15-20 years), young adults (aged 21-35 years) and adults (aged 36-100 years). All participants were enrolled to the project using electronic data capture fortified with biometric fingerprint technology for the registration and subsequent identification of participants. This generated an 11-digit participant number, which could be identified either biometrically, by scanning their ID card, or searching for their name. Where sentinel sites existed in censused districts (n = 16 kebeles), participants could be linked to their census record and cMDA record.

All faecal samples collected from a participant were provided with a six-digit sample number identifiable through scanning a QR code. This code was placed on sample pots and Kato Katz diagnostic slides to reduce human errors in labelling of samples. This also allowed for anonymised diagnostics in the laboratory. Quadruple Kato Katz slides were analysed, generated from two slides prepared per sample over two consecutive days. A minimum of six months was allowed between cMDA distribution and sentinel site parasitological monitoring to ensure any recrudescence (parasite reinfection) could be captured, signifying continued STH transmission.

To date, seven rounds of cMDA have been delivered across five years of treatment to Korke Doge. Delivery methods of cMDA were adapted during the project due to the COVID-19 pandemic, including the replacement of fixed-point distribution employed prior to the pandemic by house-to-house distribution, avoiding mass congregation of individuals within the community. Anthelmintics were distributed by the existing network of HEW, a network of care workers stationed in each kebele across Ethiopia underpinning the national healthcare system. A single dose of albendazole was offered to members of the community aged one year old and above, with syrup distributed to infants aged one to four years old. From Year 4 onwards, albendazole was offered biannually.

During cMDA, each participant was identified using their biometric fingerprint or ID card, which linked their 11-digit participant number to their treatment behaviour. Each participant was therefore categorised as either ‘missed’ or ‘contacted’ by a HEW, with the latter further categorised as ‘accepting’ or ‘refusing’ the offered drug, and subsequently ‘refusing’ or ‘swallowing’ the accepted drug, as recorded by the HEW. The eligible population for albendazole was calculated as the population aged one year old and above, excluding all pregnant women.

**DNA extraction and sequencing**

DNA extraction was performed on snips sections of worm tissue across 66 adult worm samples. This was performed using the Qiagen MagAttract magnetic beads extraction methods across all worms. DNA quality and quantity was performed through the use of gel electrophoresis and the use of Qubit quantification assay.

Paired-End Genome Libraries – Sixty-eight A. lumbricoides DNA samples were sequenced using Illumina HiSeq 2500 (www.illumina.com) short-read paired-end sequencing. DNA was quantified by UV Spec and Picogreen. A 100 ng of DNA based on picogreen quantification was used as template for NGS library preparation using the TruSeq Nano DNA Sample library prep kit. Primer-dimers in the libraries were removed by additional AMPure beads purification. Sequencing was performed to obtain a minimum genomic depth of 20X coverage for each sample. Mate-Pair Genome Libraries – Two samples were selected for mate-pair sequencing, based on the quality of the DNA preparation.. The mate-pair libraries were generated using the Nextera Mate Pair Library Prep Kit, following the gel-free method with the only modification that M-270 Streptavidin binding beads were used instead of M-280 beads. The libraries were amplified for 15 cycles given the low DNA input going into the circularization phase. The mate-pair fragment size averaged 6 kb with a range of 2–10 kb fragments.

**Variant discovery and annotation**

Raw sequence reads from all 66 samples were timed using fastqc (Patel & Jain, 2012) to remove low-quality bases and adapter sequences. Trimmed sequences were aligned to *A. lumbricoides* (AL V5) (Easton et al., 2020) reference genome. Trimmed sequence reads were aligned using BWA mem (Li & Durbin, 2009). PCR Duplicates were marked using PicardTools MarkDuplicates (McKenna et al., 2010). Variant calling was performed using GATK HaplotypeCaller in gVCF mode, retaining both variant and invariant. The individual gVCFs were merged using GATK CombineGVCFs, and joint-call cohort genotyping was performed using GATK GenotypeGVCF’s. Variant sites with on single-nucleotide polymorphisms (SNPs) were separated from indels and mixed sites (variant sites having both SNPs and indels) using GATK SelectVariants. GATK VariantFiltration was used to filter both these independently. SNPs were retained when meeting the following criteria in the filtering process; QD ≥ 2.0, FS ≤ 60.0, MQ ≥ 40.0, MQRankSum ≥ −12.5, ReadPosRankSum ≥ −8.0, SOR ≤ 3.0. Additional filtering including the Variant sites that contained wither indels or mixed sites were also retained when meeting the following; QD ≥ 2.0, FS ≤ 200.0, ReadPosRankSum ≥ −20.0, SOR ≤ 10.0.

VCFtools (Danecek et al., 2011) was used to exclude accessions with a high rate of variant sire missingness (>5% of sites calling missing genotype), subsequently removing sites where >10% of accessions had a missing genotype. This filter process formed the primary VCF file for downstream analysis. To analyse nucleotide diversity (Pi) and fixation index (*F_ST_*), a second VCF file was produced. We used the VCFtools to filter both variant and invariant sites with >80% missing variants, minimum mean read depth of 5 and maximum read depth of 500. Variant sites that were found to be significantly out of Hardy-Weinberg equilibrium (*p* < 0.001). Functional annotation of SNPs and indels in the primary analysis VCF file was performed using SnpEff (v.5.0e) (Cingolani et al., 2012) with gene annotations downloaded from WormBase ParaSite (Howe et al., 2017).

**Depth of coverage**

The depth of read coverage was calculated in 2 and 25kb windows along each chromosome using bedtools coverage (Quinlan & Hall, 2010).

**Population genomic structure and diversity**

Variants that were found at minor allele frequencies < 0.05 and excluded all variants found on sex chromosomes X1, X2, X3, X4 and X5. Additionally, we removed all variants found within repetitive regions identified using RepeatMasker (Nishimura, 2000). Finally, we removed all variants found to be in strong linkage disequilibrium. Principal component analysis was performed on the remaining 1,843,016 autosomal SNP using PLINK (Purcell et al., 2007). Genomic structure was analysed using Admixture (Meisner & Albrechtsen, 2018)with *K* values (hypothetical ancestral populations) ranging from 1 to 20, 10 fold cross-validation, standard error estimation with 500 bootstraps, this entire process was repeated 20 times with different random seeds.

The 1,843,016 autosomal variants were used to construct a neighbour joining tree. An identity-by-state distance matrix with distances expressed as genomic proportions generated by PLINK . The ape bionj algorithm (Paradis & Schliep, 2019) was performed on the resulting matrix and visualised using ggtree (Yu et al., 2017).

PIXY (Korunes & Samuk, 2021) was used to calculate the nucleotide diversity (π), fixation index (*F_ST_*) and absolute divergence (*d*_XY_), in 5 kb sliding windows with no-overlap across autosomes for each household and designated compliance dataset. Negative *F_ST_* values were corrected to 0 prior to calculation of genome wide median values for each population. Confidence intervals for these median values were calculated as the quantiles of the distribution of medians of 1000 bootstrap samples of genomic windows for each population.

**Spatial population genomics**

To understand the extent of genetic structuring and identify spatial genetic discontinuities across the endemic community. We used several related methods which are detailed below; in brief we used Bayesian clustering with BAPS v5.3 (Corander et al., 2008) to test for the presence of multiple genetic populations under Hardy-Weinberg equilibrium, principal components analysis (PCA) to visually identify major trends in the genomic structuring through ordination, fineSTRUCTURE to assess genetic structuring through differences in shared coancestry (Malinsky et al., 2018); spatial principal components analysis to assess the genetic structuring that captures patterns across spatial autocorrelation and genetic variability across space (ref).

When analysing samples that include multiple family groups alongside unrelated individuals, the removal of highly related pairs (full-siblings) allows more accurate quantification of population genetic structure and diversity (ref ref). Using PLINK *genome* function identifies pairs of individuals with a kinship coefficient greater than 0.4. When pairs of these individuals are identified, we removed whichever individual had the greatest amount of missing data, this resulted in a filtered data set of 54 individuals.

The program BAPS was used to determine the most likely number of putative genetic clusters (K) by maximising the HWE through a stochastic search algorithm. The algorithm uses sample location as a spatial prior. The module defined ‘spatial clustering of groups’ was used to examine values ranging from 1 to 20 in five independent iterations, using a reduced dataset of 50,000 random SNPs was performed for computational ease.

The programme FINESTRUCTURE was used to fit a model of population structure estimating shared co-ancestry across recombination blocks on individual’s chromosomes. First loci was phased using shapeit4 (ref), then ran FINESTRUCTURE using the unlinked model for 100,000 MCMC iterations and 20,000 tress building iterations, using a minimum of 500 SNPs and 10% of the genome for the expected maximisation estimation. The co-ancestry matrix was analysed and plotted through the tidyheatmap package in R (Mangiola & Papenfuss, 2020).

sPCA is a spatially explicit ordination approach implemented through the ADEGENET R package (Jombart, 2008). The model identifies eigenvectors that maximize both genetic variance and trends in spatial autocorrelation . This analysis identifies both “global” structures, through the presence of positive autocorrelation, indicating clinal patterns in population differentiation, as well as “ local” structures, highlighted through the presence of negative autocorrelation that occurs as genetic distance changes rapidly over small spatial scales (Jombart & Ahmed, 2011). We implemented the sPCA specific test between households in the community to assess the statistical significance of spatial structure to genetic clustering. A Gabirel graph was used to create a spatial network within sPCA and calculated *F_ST_* between identified clusters using the PIXY package (ref).
